# Germinal Center Cytokines Driven Epigenetic Control of Epstein-Barr Virus Latency Gene Expression

**DOI:** 10.1101/2024.01.02.573986

**Authors:** Yifei Liao, Jinjie Yan, Nina R. Beri, Roth G. Lisa, Cesarman Ethel, Benjamin E. Gewurz

## Abstract

Epstein-Barr virus (EBV) persistently infects 95% of adults worldwide and is associated with multiple human lymphomas that express characteristic EBV latency programs used by the virus to navigate the B-cell compartment. Upon primary infection, the EBV latency III program, comprised of six Epstein-Barr Nuclear Antigens (EBNA) and two Latent Membrane Protein (LMP) antigens, drives infected B-cells into germinal center (GC). By incompletely understood mechanisms, GC microenvironmental cues trigger the EBV genome to switch to the latency II program, comprised of EBNA1, LMP1 and LMP2A and observed in GC-derived Hodgkin lymphoma. To gain insights into pathways and epigenetic mechanisms that control EBV latency reprogramming as EBV-infected B-cells encounter microenvironmental cues, we characterized GC cytokine effects on EBV latency protein expression and on the EBV epigenome. We confirmed and extended prior studies highlighting GC cytokine effects in support of the latency II transition. The T-follicular helper cytokine interleukin 21 (IL-21), which is a major regulator of GC responses, and to a lesser extent IL-4 and IL-10, hyper-induced LMP1 expression, while repressing EBNA expression. However, follicular dendritic cell cytokines including IL-15 and IL-27 downmodulate EBNA but not LMP1 expression.

CRISPR editing highlighted that STAT3 and STAT5 were necessary for cytokine mediated EBNA silencing via epigenetic effects at the EBV genomic C promoter. By contrast, STAT3 was instead necessary for LMP1 promoter epigenetic remodeling, including gain of activating histone chromatin marks and loss of repressive polycomb repressive complex silencing marks. Thus, EBV has evolved to coopt STAT signaling to oppositely regulate the epigenetic status of key viral genomic promoters in response to GC cytokine cues.

**Author Summary:** A longstanding question has remained how Epstein-Barr virus (EBV) epigenetically switches between latency programs as it navigates the B-cell compartment. EBV uses its latency III program to stimulate newly infected B cell growth and then trafficking into secondary lymphoid tissue germinal centers (GC). In latency III, the viral C promoter stimulates expression of six Epstein-Barr nuclear antigens (EBNA) that in turn induce two latent membrane proteins (LMP). However, knowledge has remained incomplete about how GC microenvironmental cues trigger switching to latency II, where only one EBNA and two LMP are expressed, a program observed in Hodgkin lymphoma. Building on prior evidence that GC cytokines are a major cue, we systematically tested effects of cytokines secreted by GC-resident T follicular helper and follicular dendritic cells on EBV latency gene expression and on epigenetic remodeling of their promoters. This highlighted that a range of GC cytokines repress latency III EBNA, while only several support LMP1 expression, major events in the transition between the latency III and II programs. We identified key downstream roles of JAK/STAT signaling in relaying cytokine signals to the EBV epigenome, including obligatory STAT3 and 5 roles in rewiring of C and LMP promoter histone epigenetic marks.

## Introduction

Epstein-Barr virus (EBV) persistently infects >95% of adults worldwide. Although typically benign, EBV nonetheless contributes to approximately 1.5% of all human cancers.^1^ These include endemic Burkitt lymphoma (BL), Hodgkin lymphoma, natural killer/T cell lymphoma, post-transplant lymphoproliferative disease (PTLD), primary central nervous system lymphoma and diffuse large B-cell lymphoma, which typically arise from the germinal center (GC).^1–3^ EBV is also highly associated with multiple sclerosis.^4, 5^ According to the EBV GC model, EBV uses distinct combinations of latent membrane proteins (LMP) and Epstein-Barr nuclear antigens (EBNA) to expand the pool of infected B-cells, navigate the B-cell compartment and promote infected cell differentiation into memory B-cells, the reservoir for lifelong infection.^6^ Across these latency programs, ∼80 viral lytic antigens are largely silenced by epigenetic mechanisms.

The EBV genome is epigenetically programmed upon B cell infection.^7–10^ While EBV genomic DNA is epigenetically naïve in viral particles, it is rapidly chromatinized as incoming viral genomes reach the infected cell nucleus.^7, 11^ Histone epigenetic marks, DNA methylation and three dimensional EBV genomic architecture then serve as major regulators of EBV gene expression. Much remains to be learned about host cell transcription factors and their upstream pathways in control of EBV epigenomic programming. The viral W promoter (Wp) drives an initial burst of EBNA expression, in particular EBNA2 and EBNA-LP, which highly upregulate MYC and other key B-cell targets.^12–19^ Infected cells then transition to the latency IIb program, in which the EBV genomic C promoter (Cp) drives expression of a transcript encoding EBNAs 1, 2, 3A, 3B, 3C and LP, whose messages are subsequently spliced. Shortly thereafter, EBNA2 activates the latent membrane promoters, driving expression also of LMP1 and LMP2A, culminating in the latency III program.^3^ If left unchecked, the transforming latency III program converts B-cells into immortalized lymphoblastoid cell lines (LCL), a key model for PTLD and AIDS-associated immunoblastic lymphomas.^1, 9, 20^

Latency III drives cells into GC, where immune pressure together with incompletely understood mechanisms are believed to drive the transition to the EBV latency II program, comprised of EBNA1, LMP1 and 2A.^9^ EBNA1 expression is driven by the viral genome Q promoter (Qp) in latency II. Much remains to be understood about the precise GC signals and their downstream epigenetic mechanisms that culminate in Cp silencing, while instead supporting LMP expression in the absence of EBNA2 transcription activation. Upon memory B-cell differentiation, epigenetic mechanisms likely including DNA methylation and polycomb repressor complex 1 silence the LMP promoters to enable progression latency I program, where EBNA1 is the only EBV- encoded protein expressed.^8, 21^

The GC is a dynamic secondary lymphoid tissue microstructure, where T follicular helper (Tfh) and follicular dendritic cells (FDC) together with antigens drive B-cell responses.^22, 23^ Tfh cytokines, including IL-2, 4, 10, and 21, together with the FDC derived cytokines IL-6, 15 and 27, are critical for GC establishment and maintenance, as well as for GC B-cell fate.^23–26^ Cytokines bind to plasma membrane B-cell receptors to activate Janus kinase (JAK) or Tyrosine kinase 2 (TYK2), which phosphorylate specific signal transducer and activator of transcription (STAT) family proteins.

Phosphorylation drives STAT dimerization via reciprocal SH2 domain–phosphotyrosine interactions and nuclear translocation to enable target gene regulation (**Fig. S1A**).^27–29^ IL-21 decreases EBNA2 expression in latency III B cells^30, 31^, suggesting a potential GC cytokine role in driving the transition from latency III to II. Moreover, IL-4, 10, and 21 each de-repress LMP1 expression in newly infected cells and in latency I Burkitt and natural killer (NK) lymphoma cells^30–36^, further suggesting roles in support of latency II. IL-15 also drives NK and T-cell responses against EBV transformed peripheral blood B-cells^37, 38^, potentially suggesting that it may enhance immune pressure against latency III B-cells within the GC. However, much remains to be learned about the mechanisms by which cytokines secreted by Tfh and FDC alter the EBV epigenome to repress EBNA but instead support LMP expression.

To gain insights into mechanisms by which GC cytokines alter EBV latency gene expression and the viral epigenome, we systematically screened effects of Tfh and FDC cytokines on EBV latency gene expression. Tfh cytokines, including IL-4, 10 and 21, each upregulated LMP1 but downregulated EBNA2 and 3 levels in B cells with latency III. By contrast, the key FDC cytokine IL-15 diminished Cp driven EBNA expression but did not significantly alter LMP1 levels. CRISPR analysis identified that STAT3 and to a lesser extent STAT5 was critical for these cytokine effects on EBNA and LMP1 expression. Taken together, our results highlight GC cytokines driven STAT3 and 5 remodeling of the EBV epigenome to support the latency III to latency II program transition.

## Results

### GC cytokines support the latency II transition

To systematically characterize GC cytokine effects on EBV latency gene expression, we incubated the LCL GM12878 with a panel of Tfh-derived cytokines, IL-2, IL-4, IL-10 or IL-21. In parallel, we incubated GM12878 with the FDC-derived cytokines IL-6, IL-15 or IL-27 for 0, 2, 4 or 6 days (**Fig. 1A, Fig. S1A**). While it is not known how long EBV+ B-cells reside within the GC, it is likely that they remain present for at least several days, in order to proliferate and differentiate into memory B cells, and GC structures themselves persist for weeks to months. Cytokine effects on EBV latency programs were defined by immunoblot for EBNA2, EBNA3C and LMP1, since this panel of EBV oncoproteins can be used to assign the latency program. Interestingly, most of these cytokines reduced EBNA2 and 3C expression, though results were the most pronounced for IL-21, which rapidly and robustly impaired EBNA2/3C expression (**Fig. 1B**). By contrast, IL-10 and IL-21 upregulated LMP1 expression within 2 days of treatment (**Fig. 1B**). Similar effects were observed in a second LCL, GM12881 (**Fig. S1B**). IL-21 also suppressed EBNA2 and upregulated LMP1 in latency III Jijoye Burkitt cells (**Fig. S1C**), suggesting generalizable effects on the latency III program. Consistent with prior reports, IL-21 did not hyper-induce LMP2A expression in either GM12878 or Kem III LCLs, indicating that IL-21 may fail to induce recruitment of an activator to the LMP2 promoter or to instead dismiss a repressor (**Fig. S1D**).

**Figure 1.**
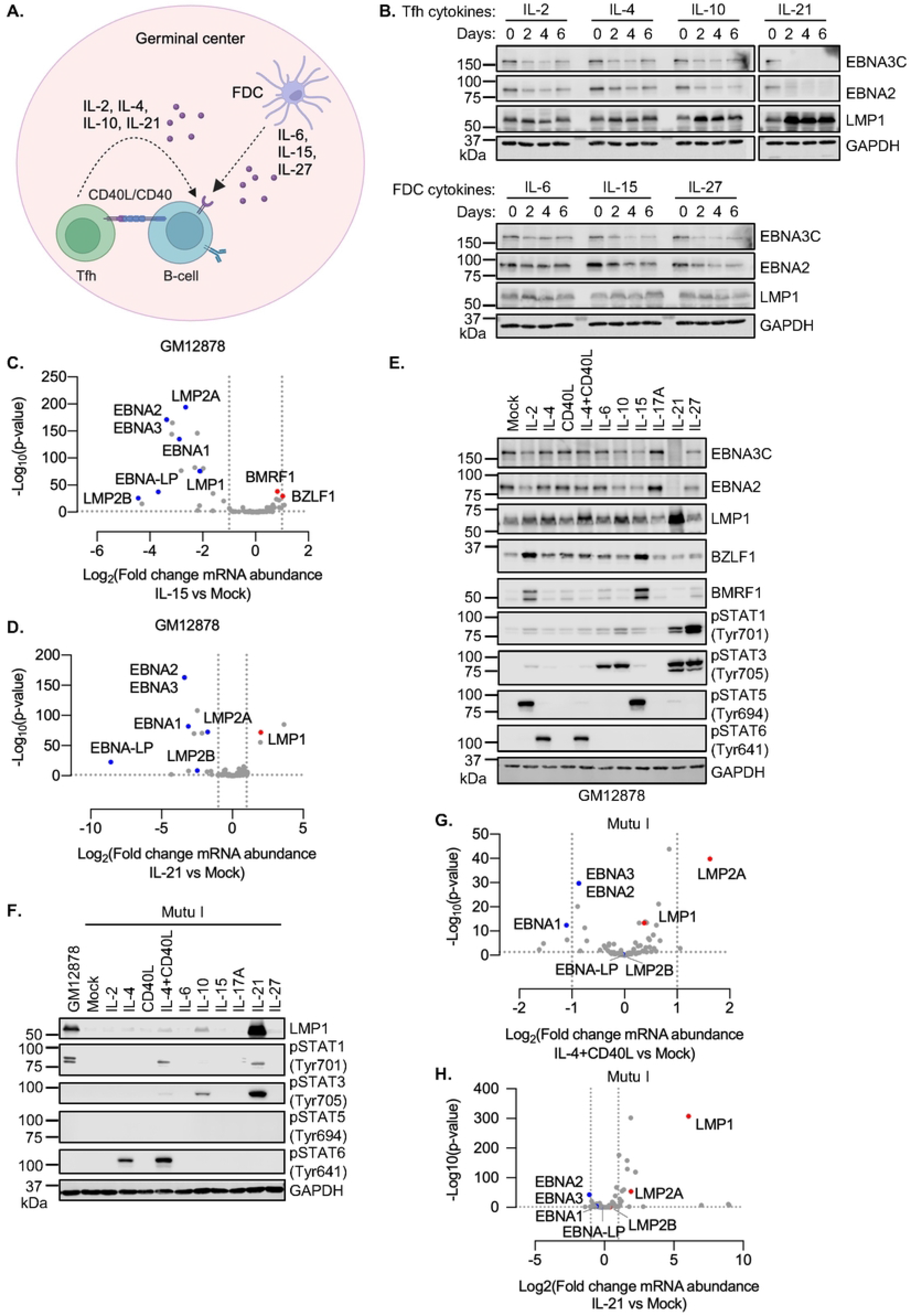
GC cytokines support the transition to EBV latency II. (**A**) GC schematic, illustrating key T follicular helper cell (Tfh) and follicular dendritic cell (FDC) secreted cytokines and CD40 ligand (CD40L) that signal to GC B-cells. (**B**) Immunoblot analysis of whole cell lysates (WCL) from GM12878 treated with the indicated cytokines for two, four, or six days. (**C**) Volcano plots of EBV gene expression from n = 3 replicates of GM12878 stimulated by IL-15 vs. mock-stimulated for 6 days. (**D**) Volcano plots of EBV gene expression from n = 3 replicates of GM12878 stimulated by IL-21 vs. mock-stimulated for 6 days. (**E**) Immunoblot analysis of WCL from GM12878 cells treated with the indicated cytokines for six days. (**F**) Immunoblot analysis of WCL from Mutu I cells treated with the indicated cytokine for one day or from GM12878 for comparison. (**G**) Volcano plot of EBV gene expression from n = 3 replicates of Mutu I stimulated by IL- 4+CD40L vs. mock-stimulated for one day. (**H**) Volcano plot of EBV gene expression from n = 3 replicates of Mutu I stimulated by IL-21 vs. mock-stimulated for one day. All cytokines were used at 100 ng/ml and 50 ng/ml in GM12878 vs Mutu I, respectively, and were refreshed every two days. Immunoblots are representative of n = 3 replicates.

We next performed RNA-seq analysis to systematically characterize IL-15 and IL-21 effects on EBV genome wide expression, as representative of FDC vs Tfh cytokine signaling, respectively. After six days of treatment, IL-15 significantly decreased expression of multiple latency III genes, including EBNA2, EBNA3, EBNA-LP, LMP1 and LMP2A, but increased expression of a subset of lytic cycle genes, including immediate early BZLF1 and early BMRF1 (**Fig. 1C, Table S1**), suggestive of an abortive lytic cycle. Instead, IL-21 significantly increased abundance of LMP1 mRNA but decreased abundances of EBNA2, EBNA3, EBNA-LP, and LMP2 mRNAs (**Fig. 1D, Table S1**). Consistent with effects on EBNA2 and LMP1 expression, IL-15 downregulated the EBNA2 target gene CD300A, while IL-21 upregulated levels of the LMP1/NF-κB target ICAM-1 and downmodulated CD300A^21, 39, 40^ (**Fig. S1E-F, Table S2**).

GC cytokines alter B-cell gene expression patterns via multiple effectors, including distinct JAK/STAT pathways. As expected, the panel of cytokines differentially activated STATs, including STAT5 activation by IL-2 and IL-15 versus STAT6 activation by IL-4 versus STAT3 activation by IL-6, IL-10, IL-21 and IL-27, as judged by immunoblot for well characterized phosphorylation marks of STAT activation^26, 27^ (**Fig. 1E, Fig. S1B-C**). Consistent with our RNA-seq analyses, IL-15 de-repressed BZLF1 and BMRF1 expression at the protein level, as did IL-2 (**Fig. 1E**), suggesting that it induces an abortive lytic cycle in at least a subset of cells. Notably, these two cytokines share receptor beta and gamma chain subunits, which are transmembrane proteins that activate downstream pathways, including JAK/STAT.^41^

We next asked the extent to which Tfh and FDC cues can alter EBV latency gene expression within the latency I B-cell context. While several Tfh signals, including IL- 4+CD40L, IL-10 or IL-21 can each de-repress LMP1 expression in B-cells with the latency I program^30–34^, it has remained unknown the extent to which other GC microenvironmental cues more broadly alter EBV latency gene expression within latency I. To gain insights, we treated latency I Mutu I and Kem I Burkitt cells with a panel of Tfh and FDC cytokines, as there is no primary human B-cell latency I models currently available. IL-21 strongly activated STAT3, as judged by tyrosine 705 phosphorylation, and robustly de-repressed LMP1 expression in Mutu I and Kem I (**Fig. 1F, S2A-B**). By contrast, IL-4+CD40L or IL-10 treatment also induced STAT3 phosphorylation and LMP1 expression, albeit to a lesser extent (**Fig. 1F**). This did not appear to be a full transition to the latency II program, as neither IL-10 nor IL-21 induced LMP2A to an appreciable degree in Mutu I or Kem I (**Fig. S2C**). Differences between GC cytokine STAT activation in the latency I vs III context may relate to altered expression of receptors versus negative regulators of JAK/STAT signaling.

To then systematically analyze latency I B-cell responses to the Tfh signals IL-4+CD40L vs IL-21, we performed RNA-seq on Mutu I that were mock-stimulated or stimulated by these Tfh cues for 1 day. This early timepoint was chosen since we observed robust effects on LMP1 de-repression by that early timepoint, and as we observed reduced Mutu I viability with longer treatments. Consistent with our immunoblot analysis, IL- 4+CD40L only modestly increased LMP1 expression, whereas IL-21 strongly induced LMP1 (**Fig. 1G-H**). Notably, these stimuli did not significantly de-repress expression of EBNA or mildly increased LMP2 mRNAs, suggesting a specific effect at the level of the LMP1 promoter.

Analysis of Mutu I host transcriptome responses to either IL-4+CD40L or IL-21 treatment highlighted upregulation of multiple LMP1 target genes^42^, including mRNAs encoding the NF-κB subunits RelB and p100/52 (encoded by NFKB2), ICAM-1 and IRF4 (**Fig. S2D-E**). The NF-κB pathway signaling pathway and EBV infection were amongst the pathways most highly enriched by either cytokine treatment. While direct effects of the cytokines themselves may account for a subset of these changes, we note that IL-21 is not a strong inducer of NF-κB signaling, suggesting that de-repressed LMP1 may be an important mediator of the observed host transcriptomic changes.

#### STAT3 and STAT5 mediate GC cytokine effects on the EBV latency III program

We next investigated effects of chemical or CRISPR JAK/STAT blockade to gain further insight into specific STAT roles in modulation of EBV latency oncogene expression downstream of IL-15 and IL-21. First, to broadly characterize JAK/STAT roles in LMP and EBNA expression, we treated latency III GM12878 and Jijoye cells with IL-15 or IL- 21, in the absence or presence of the pan-JAK ATP-competitive inhibitor CAS 457081- 03-7 (also referred to as JAK inhibitor I or JAKi). On-target JAKi effects were confirmed by immunoblot analysis of STAT3 and STAT5 phosphorylation, which demonstrated loss of STAT5 Tyrosine 694 and downmodulation of STAT3 Tyrosine 705 phosphorylation in IL-15 and IL-21 treated cells, respectively (**Fig. 2A-B**). JAKi impaired IL-15 downmodulation of EBNA2 and 3C expression (**Fig. 2A-B**). Likewise, JAKi treatment partially impaired IL-21 suppression of EBNA2 and EBNA3C expression and reduced the extent to which IL-21 hyper-induced LMP1 (**Fig. 2A-B**). Incomplete blockade of IL-21 driven STAT3 phosphorylation may explain the comparatively milder JAKi effects on IL-21 than on IL15 regulation of latency III expression.

**Figure 2.**
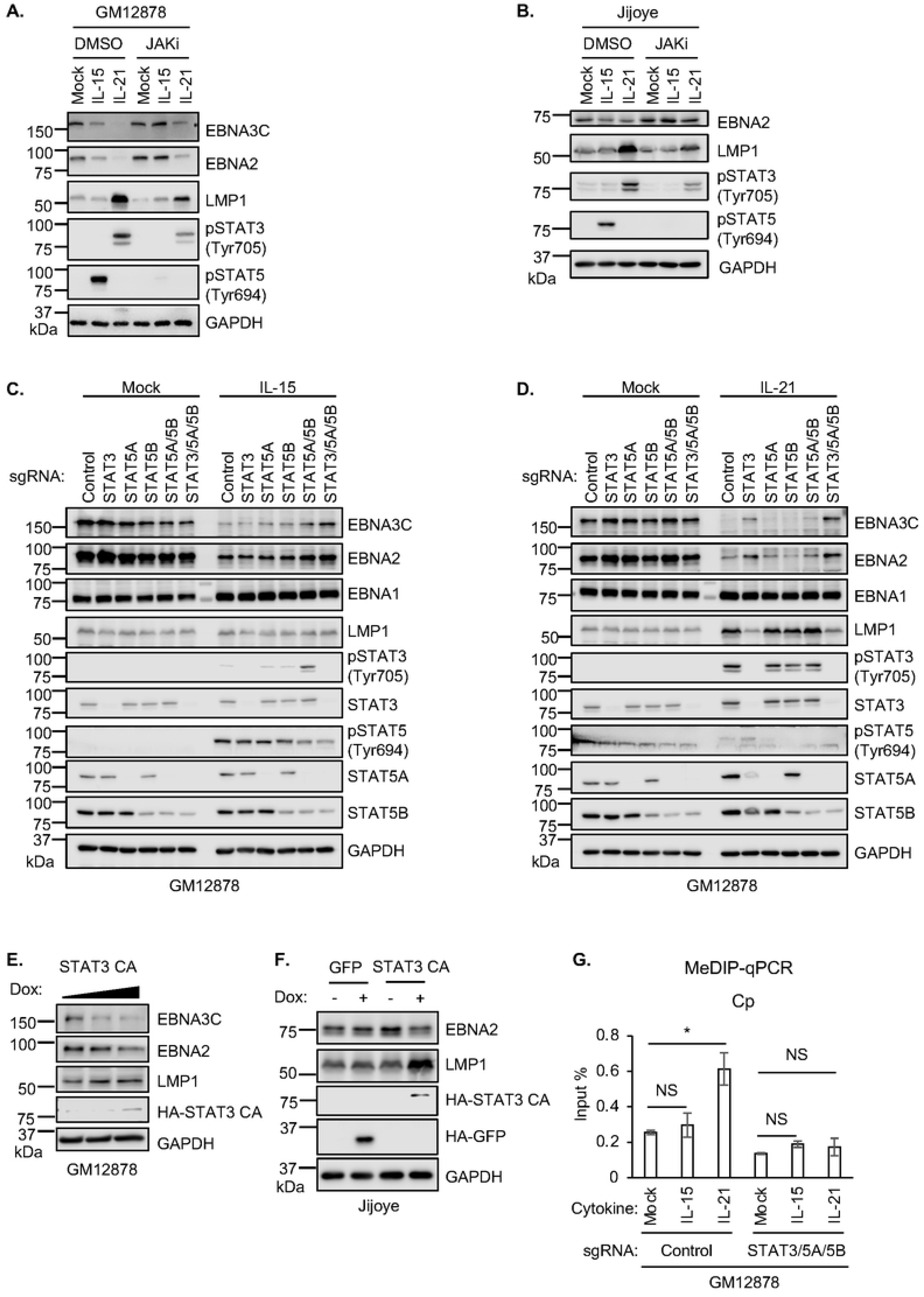
STAT3 and 5 roles in IL-15 and IL-21 driven EBV latency III gene regulation. (**A-B**) Immunoblot analysis of WCL from GM12878 (A) or Jijoye (B) pre-treated with DMSO vehicle or JAK inhibitor CAS 457081-03-7 (JAKi, 200 ng/ml) for one hour, followed by treatment with IL-15 or IL-21 for six days. (**C-D**) Immunoblot analyses of WCL from GM12878 cells expressing control sgRNA versus sgRNA targeting the indicated STAT3 and/or STAT5 genes, mock treated or treated with IL-15 (C) or IL-21 (D) for six days. (**E-F**) Immunoblot analysis of WCL from GM12878 (E) or Jijoye (F) induced for GFP or constitutive activated STAT3 (STAT-CA) by 0.5 or 1 µg/ml doxycycline (Dox). (**G**) Methylated DNA immunoprecipitation and quantitative PCR (MeDIP-qPCR) analysis of GM12878 with control or STAT3/5A/5B sgRNA expression, mock treated or treated with IL-15 or IL-21 for six days. Shown is mean ± standard deviation (SD) from n = 3 replicates of Cp qPCR signal. \**p* < 0.05; NS: not significant. IL-15 and IL-21 were used at 100 ng/ml throughout. Immunoblots are representative of n = 3 replicates.

To examine individual STAT transcription factor roles downstream of IL-15 or IL-21, we next used CRISPR/Cas9 editing. Since IL-15 and IL-21 most robustly induced STAT5 and STAT3 phosphorylation (**Fig. 1E**), we tested effects of CRISPR depletion of STAT3, of STAT5A or STAT5B isoforms^43^, or of combinations thereof, given their potentially redundant roles. IL-15 repression of EBNA2 or 3C was not significantly perturbed by depletion of STAT3, STAT5A or STAT5B alone. However, concurrent GM12878 and Jijoye STAT5A/5B depletion impaired repression of EBNA2 and 3C by IL-15 and to a lesser extent by IL-21 (**Fig. 2C-D, S3A**). However, concurrent CRISPR depletion of STAT3, STAT5A and STAT5B more strongly impaired EBNA3C repression by IL-15 (**Fig. 2C**), suggestive of a partially redundant STAT3 and 5 roles, likely at the EBV C promoter.

STAT3 depletion was sufficient to block IL-21 driven LMP1 hyper-induction and impaired IL-21 driven EBNA2/EBNA3C repression (**Fig. 2D**). Nonetheless, combined STAT3/5A/5B editing more strongly impaired EBNA2 and EBNA3C repression by IL-21 (**Fig. 2D**). Despite robust STAT1 activation by IL-21 and to a lesser extent by IL-10 (**Fig. 1E**), CRISPR STAT1 depletion did not alter IL-21 or IL-10 effects on EBNA or LMP1 expression (**Fig. S3B-C**). STAT3 KO impaired EBNA2/3C repression and LMP1 hyper-induction downstream of IL-10 (**Fig. S3B-C**). Taken together, these results suggest that STAT3 and 5 have partially redundant roles in cytokine mediated EBNA2/3C repression, perhaps through the action of STAT3/5 heterodimers, whereas STAT3 is a major driver of IL-21 driven LMP1 hyper-induction, with relevance to latency III to II reprogramming in the GC microenvironment.

Since IL-15 and IL-21 upregulated the host transcriptional repressor BCL6 (**Fig. S1E-F**) which plays major roles in GC B-cell biology and is critical for GC formation, we tested BCL6 roles in cytokine driven EBV latency gene expression. However, BCL6 CRISPR KO did not appreciably alter IL-21 effects on EBNA2 or LMP1 abundance (**Fig. S4A**).

BCL6 KO also did not affect IL-21 effects on LCL plasma membrane CD300A or ICAM- 1, which are targets of EBNA2 and LMP1, respectively (**Fig. S4B**). By contrast, expression of a constitutively active STAT3 allele with A662C and N664C point mutations^44^ diminished EBNA2 and increased LMP1 abundance in GM12878 and Jijoye B-cells (**Fig. 2E-F**), further suggesting that STAT3 plays a critical but opposite role in EBNA2/3 vs LMP1 regulation. These observations are consistent with a model in which GC cytokine signaling culminates in assembly of STAT3/5-containing transcriptional repressor complexes at the EBV genomic C promoter, but instead triggers formation of a STAT3 homodimer containing activator complex at the LMP1 promoter.

DNA methylation is critical for suppression of Cp driven EBNA expression in B-cells with latency I, and presumably also in latency II^21, 45–48^. We therefore used methylation DNA immunoprecipitation (MeDIP) and qPCR to characterize IL-15 versus IL-21 effects on LCL Cp DNA methylation levels. Notably, IL-21, but not IL-15 significantly increased C, LMP1 and LMP2A promoter methylation levels, and STAT3/5A/5B depletion reversed this effect (**Fig. 2G, Fig. S4C**). These results indicate that STAT3/5 promote cross-talk between IL-21, C and LMP promoter DNA methylation.

### STAT and DNA methylation roles in latency I LMP1 de-repression by GC cytokines

To gain insights into JAK/STAT roles in GC cytokine triggered LMP1 de-repression in latency I B-cells, we treated Mutu I or Kem I Burkitt cells with IL-10, IL-21 or IL-4 together with CD40 ligand, in the absence or presence of JAK inhibition. We tested these GC stimuli since each hyper-induced LMP1 and robustly induced STAT phosphorylation in latency I cells and had previously been reported to de-repress LMP1 expression from latency I^30–36^ (**Fig. 1F**). JAKi treatment strongly impaired LMP1 upregulation by each of these stimuli (**Fig. 3A, S5A**), consistent with a key JAK/STAT role in epigenetic regulation at the level of the LMP1 promoter.

**Figure 3.**
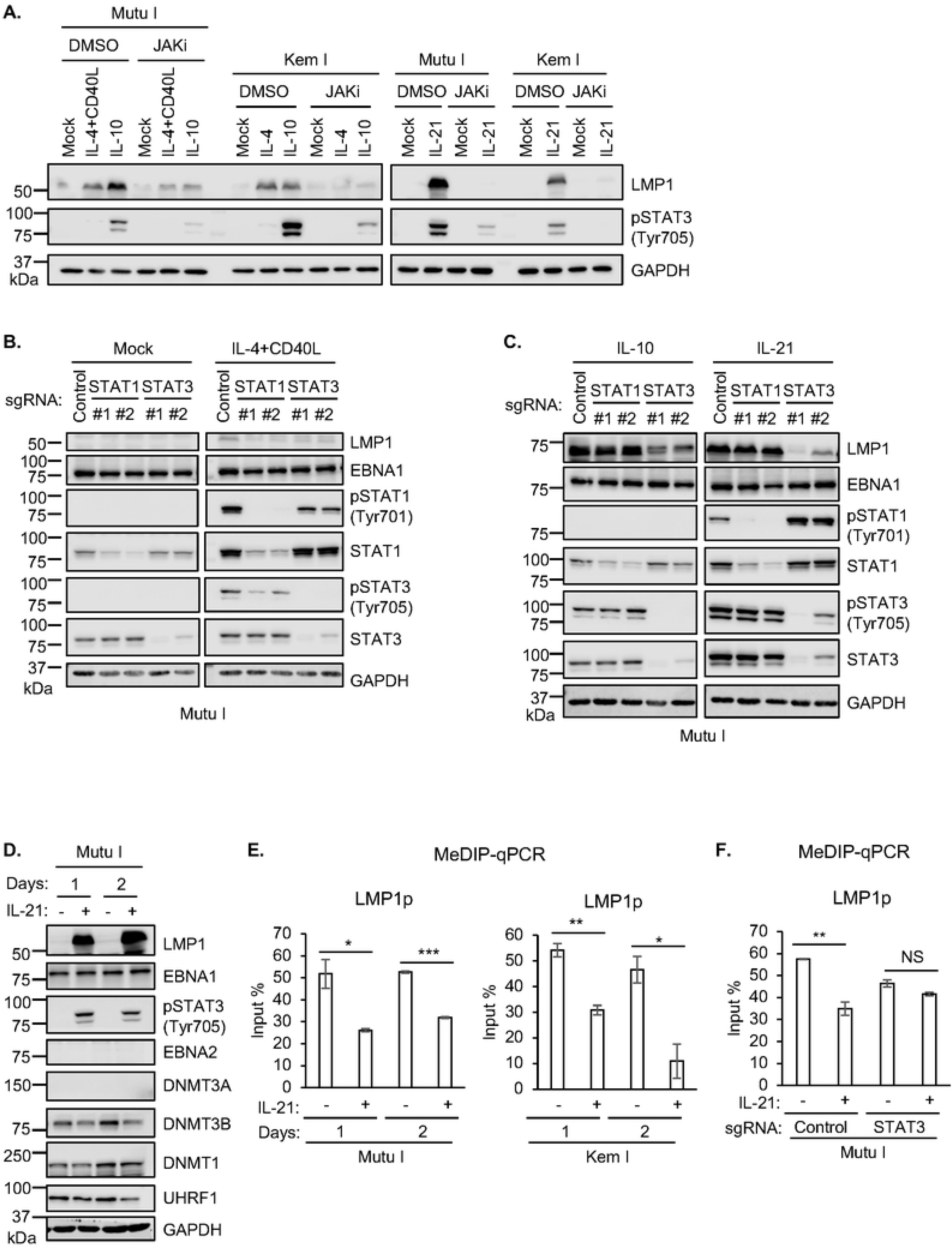
STAT3 roles in GC cytokine mediated LMP1 de-repression in latency I B- cells. (**A**) Immunoblot analysis of WCL from latency I Mutu I or Kem I B cells pre-treated with DMSO or JAKi (200 ng/ml) for one hour, followed by treatment with the indicated cytokines for one day. (**B**) Immunoblot analysis of WCL from Mutu I expressing control sgRNA or sgRNA targeting the indicated STAT transcription factor gene, mock treated or treated with IL-4+CD40L for one day. (**C**) Immunoblot analysis of WCL from Mutu I expressing control sgRNA or sgRNA targeting the indicated STAT transcription factor gene, treated with IL-10 or IL-21 for one day. (**D**) Immunoblot analysis of WCL from Mutu I mock treated or treated with IL-21 for one or two days. (**E**) MeDIP q-PCR analysis of LMP1 promoter methylation in Mutu I (left) or Kem I (right) mock treated or treated with IL-21 for one or two days. Shown are mean ± SD values of % input from n = 3 replicates. (**F**) MeDIP-qPCR analysis of LMP1p in Mutu I with control or *STAT3* targeting sgRNA, mock treated or IL-21 treated for one day. \**p* < 0.05; \*\**p* < 0.01; \*\*\**p* < 0.001. Cytokines were used at 50 ng/ml. Blots are representative of n = 3 replicates.

To next gain mechanistic insights into specific STAT roles, we next tested effects of CRISPR depletion of STAT transcription factors that were highly phosphorylated in response to these GC stimuli (**Fig. 1F**). Depletion of either STAT1 or STAT3 blunted LMP1 de-repression by IL-4+CD40L stimulation or even by IL-4 alone (**Fig. 3B, S5B**). By contrast, depletion of STAT3, but not STAT1, impaired IL-10 and IL-21 mediated LMP1 de-repression in Mutu I and Kem I cells (**Fig. 3C, S5B**). Thus, STAT1/3 heterodimers may be important for IL-4 driven LMP1 de-repression, whereas distinct STAT3 heterodimers or homodimers may mediate LMP1 de-repression downstream of IL-10 and IL-21. Consistent with the latter hypothesis, induction of the constitutively active STAT3 allele was sufficient to de-repress LMP1 expression in Mutu I (**Fig. S5C**). Likewise, STAT3 and to a somewhat lesser extent STAT6 over-expression enhanced LMP1 de-repression in response to cytokine treatment (**Fig. S5D**).

To gain further insights into cytokine cross-talk with EBV-genomic CpG methylation, we next analyzed IL-21 effects on the abundance of DNA methyltransferase machinery. IL- 21 downregulated expression of the *de novo* CpG methylation writer DNMT3B, whose expression counteracts latency III gene expression.^21^ Likewise, IL-21 downmodulated UHRF1 expression, which is important for maintenance of EBV genomic methylation marks, together with DNMT1.^21^ Therefore, to further characterize IL-21 effects on CpG methylation of key EBV genomic promoters, we performed MeDIP-qPCR analysis on Mutu I or Kem I Burkitt cells treated with IL-21 for 1 or 2 days. Interestingly, IL-21 downmodulated the high level of DNA methylation at the LMP1 and C promoters, but not at the LMP2 promoter in either Mutu I or Kem I (**Fig 3E, S5E**). Additional epigenetic marks may maintain Cp and LMP2p silencing upon IL-21 stimulation in the latency I context, including those driven by STAT-containing repressive complexes. In support of a STAT3 role in modulation of LMP1p methylation downstream of IL-21, we did not observe diminished LMP1p methylation levels in STAT3 depleted Mutu I cells upon IL- 21 treatment (**Fig. 3F**).

### GC cytokine effects on LMP1 promoter histone epigenetic marks

In addition to DNA methylation, histone epigenetic marks strongly contribute to EBV latency gene expression.^14, 21, 49–56^ We therefore next profiled GC cytokine effects on the LMP1 promoter. Since previous studies identified three LMP1 promoter sites occupied by STAT factors^57^ (**Fig. 4A**), we performed chromatin immunoprecipitation (ChIP) and qPCR analyses in LCLs mock treated or treated with FDC-derived IL-15 or Tfh-derived IL-21. IL-21 increased STAT3 occupancy at the S3 site, located at approximately 600 base pairs (bp) upstream of LMP1p, and to a lesser extent at the S2 and S1 sites, located at approximately 500 and 100 bp upstream of LMP1p (**Fig. 4B**). Interestingly,

**Figure 4.**
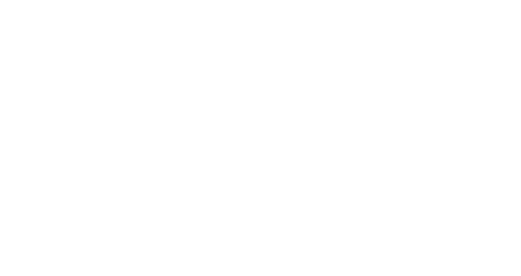
STAT 3 and 5 roles in GC cytokine mediated latency III LMP1 promoter epigenetic remodeling. (**A**) Schematic diagram of LMP1 promoter STAT binding sites, S1, S2 and S3.^57^ (**B-C**) Chromatin immunoprecipitation (ChIP) qPCR analysis of STAT3 (B) or STAT5 (C) LMP1 promoter occupancy in GM12878 mock treated or treated with IL-15 or IL-21 for six days. (**D-E**) ChIP-qPCR analysis of LMP1 promoter H3K27Ac (D) or H2AK119Ub (E) epigenetic mark abundances in GM12878 expressing control versus STAT3/5A/5B targeting sgRNA, mock treated or treated with IL-15 or IL-21 for six days. (**F-G**) ChIP-qPCR analysis of LMP2 promoter H3K27Ac (F) or H2AK119Ub (G) abundances in GM12878 expressing control versus STAT3/5A/5B targeting sgRNA, mock treated or treated with IL-15 or IL-21 for six days. (**H-I**) ChIP-qPCR analysis of LMP1 promoter H3K27Ac (H) or H2AK119Ub (I) abundances in Mutu I expressing control sgRNA versus STAT3 targeting sgRNA, mock treated or treated with IL-21 for one day. All ChIP results are presented as % input mean ± SD from n = 3 replicates. \**p* < 0.05; \*\**p* < 0.01; \*\*\**p* < 0.001.

IL-15 instead downmodulated STAT3 occupancy at S2 and S3, consistent with the observation that it does not hyper-induce LMP1 in latency III. By contrast, IL-15 but not IL-21 significantly increased STAT5 occupancy at S1-S3 (**Fig. 4C**). These results further support the hypothesis that IL-21 driven STAT3, and potentially STAT3 homodimers, are major drivers of LMP1 hyper-induction.

To characterize STAT roles in LMP1 promoter histone epigenetic regulation, we next performed ChIP-qPCR analysis in control versus CRISPR-edited LCLs. In control LCLs, IL-15 and to a greater extent IL-21 increased LMP1p histone 3 lysine 27 acetylation (H2K27Ac), a mark which correlates with promoter activation. By contrast, IL-15 and IL- 21 failed to upregulate LMPp H3K27Ac level in LCLs depleted for STAT3, STAT5A and STAT5B (**Fig. 4D**). Since we recently found a role for the polycomb repressive complex (PRC1) I histone 2A lysine 119 ubiquitin (H2AK119Ub) mark in repression of LMP1 expression^21^, we next examined GC cytokine and STAT roles on LMP1p H2AK119Ub levels in LCLs. IL-15 and to a greater extent IL-21 significantly diminished H2AK199Ub abundance in control, but not STAT3/5A/5B KO GM12878 (**Fig. 4E**). Despite lack of appreciable LMP2A hyper-induction, IL-21 nonetheless increased H3K27Ac levels in GM12878 control and STAT3/5A/5B edited LCLs (**Fig. 4F**). Interestingly, IL-21 hyper-induced H2Ak119Ub repressive marks at LMP2p in both control and STAT3/5A/5B edited cells. Given PRC1 roles in repression of LMP expression, this result suggests a potential mechanism by which LMP1 but not LMP2A is hyper-induced in IL-21 treated B-cells, and are consistent with a model in which STAT3/5 occupy LMP1 but not LMP2 promoter sites.

We did not observe decreases in the repressive histone 3 lysine lysine 9 dimethyl (H3K9me2) or trimethyl (H3K9me3) marks with either IL-15 or IL-21 treatment in control or STAT KO LCLs (**Fig. S6A-B**). However, repressive histone 3 lysine 27 trimethyl (H3K27me3) repressive marks increased somewhat upon IL-15 or IL-21 treatment in STAT3/5A/5B triple edited LCLs (**Fig S6C**). Similar effects were observed at the LMP2A promoter, though IL-21 increased the repressive H3K9me3 mark in both control and STAT3/5A/5B edited cells (**Fig. S6D-F**), potentially contributing to the lack of IL-21 driven LMP2A hyper-induction.

We next characterized IL-21 epigenetic effects on the LMP1 promoter in latency I B-cells, given the observation that IL-21 strongly activates STAT3 phosphorylation and de- represses LMP1 expression, whereas other GC cytokine stimuli did so comparatively weakly. As anticipated, IL-21 significantly increased H3K27Ac at the LMP1 promoter in Mutu I cells. Interestingly, STAT3 was necessary for this IL-21 driven epigenetic remodeling, as STAT3 depletion prevented IL-21 driven H3K27Ac activating mark at the LMP1 promoter (**Fig. 4H**). Similarly, IL-21 significantly diminished the repressive H2AK119Ub and H3K9me2 marks at the LMP1 promoter in a STAT3 dependent manner (**Fig. 4I** **and S7A**). By contrast, IL-21 did not significantly alter repressive H3K9me3 or H3K27me3 marks at the latency I LMP1 promoter (**Fig**. **S7B**).

Interestingly, IL-21 did not significantly alter activating or repressive histone marks at the Mutu I LMP2A promoter (**Fig. S7C-F**). These results indicate that the absence of STAT3 signaling is important for silencing LMP1 expression in latency I, with relevance to the transition from latency II to latency I.

### GC cytokines remodel epigenetic status of C promoter

Multiple GC cytokines repressed latency III EBNA expression, suggestive of epigenetic effects at the level of Cp, which drives the large EBV transcript encoding all six EBNAs. We performed ChIP to characterize how IL-15 and IL-21 alter STAT3 versus STAT5 occupancy at two predicted STAT binding sites using PROMO online tool^58, 59^, located at 300 and 400 bp upstream of Cp (**Fig. 5A**). Consistent with our observation that IL-15 and IL-21 predominantly activated STAT5 versus STAT3 in latency III cells, respectively (**Fig. 1B, Fig. S1B-C**), IL-21 but not IL-15 significantly upregulated STAT3 occupancy at both the S1 and S2 sites upstream of Cp (**Fig. 5B**). Conversely, IL-15 significantly induced STAT5 occupancy at both S1 and S2, whereas IL-21 weakly induced STAT5 binding to S2 (**Fig. 5C**). Taken together with our CRISPR and immunoblot analyses, these data are compatible with a model in which a STAT5 or STAT3 homodimer and to a lesser extent a STAT3/5 heterodimer are critical for IL-15 or IL-21 mediated Cp repression.

**Figure 5.**
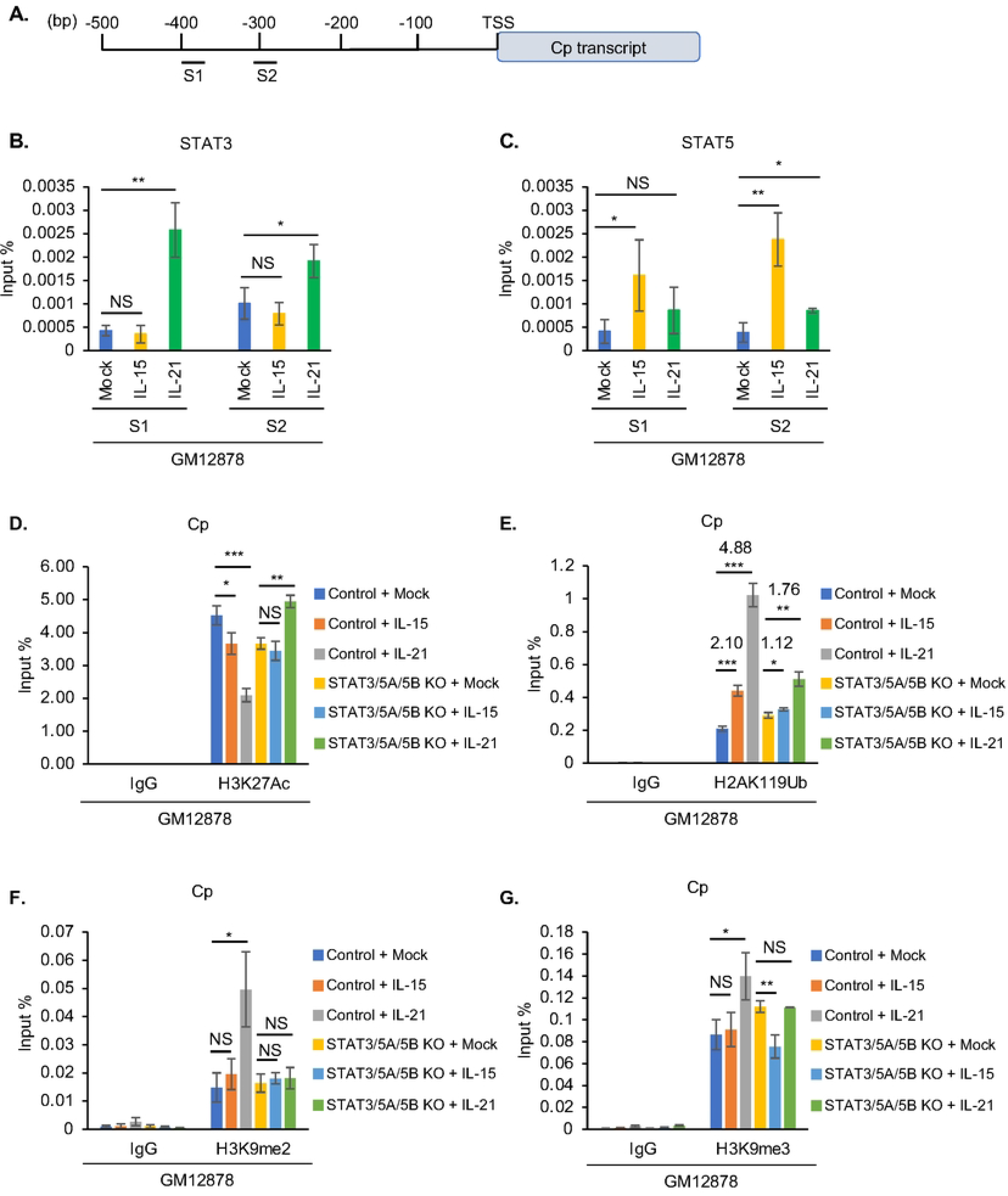
STAT3 and 5 roles in GC cytokine mediated C promoter epigenetic remodeling. (**A**) Schematic diagram of PROMO^58, 59^ predicted STAT binding sites on C promoter. (**B-C**) ChIP-qPCR analysis of STAT3 (B) or STAT5 (C) C promoter occupancy in GM12878 mock treated or treated with IL-15 or IL-21 for six days. (**D-G**) ChIP-qPCR analysis of Cp H3K27Ac (D), H2AK119Ub (E), H3K9me2 (F) and H3K9me3 (G) abundances in GM12878 expressing control versus STAT3/5A/5B targeting sgRNA, mock treated or treated with IL-15 or IL-21 for six days. All ChIP results are presented as % input mean ± SD from n = 3 replicates. \**p* < 0.05; \*\**p* < 0.01; \*\*\**p* < 0.001.

At the epigenetic level, ChIP-qPCR assays highlighted that IL-21 more strongly reduced H3K27Ac marks at Cp than IL-15. Consistent with key STAT3 and 5 roles in GC cytokine driven epigenetic remodeling at Cp, CRISPR editing of STAT3/5A/5B blocked H3K27Ac loss at Cp in GM12878 stimulated by either IL-15 or IL-21 (**Fig. 5D**). Similarly, IL-15 and to a greater extent IL-21 increased H2AK119Ub repressive marks at Cp, and CRISPR STAT3/5A/5B editing blunted cytokine-driven H2AK119Ub deposition (**Fig. 5E**). Likewise, IL-21 but not IL-15 significantly increased deposition of the H3K9me2 and H3K9me3 repressive marks at Cp, and this increase was blunted by STAT3/5 editing (**Fig. 5F-G**). Interestingly, neither IL-15 nor IL-21 increased repressive H3K27me3 marks at Cp, arguing against PRC2 roles in their repression of Cp (**Fig. S8A**). By comparison, Cp is silenced in latency I, and likely related to that, we observed relatively small differences in the Cp epigenetic status in control or STAT3 edited Mutu I at rest or following IL-21 treatment (**Fig. S8B-F**). These results are consistent with a model in which STAT3 nucleate transcription co-activator complexes at LMP1p but STAT3 and 5 mediates repressive complexes at Cp in latency III B-cells, and that latency I cells maintain the ability to respond to STAT-dependent epigenetic remodeling at LMP1p.

### JAK/STAT signaling roles in newly infected B-cell latency gene expression and transformation

JAK/STAT signaling contributes to EBV latency gene expression in newly infected primary human B-cells^60, 61^, though to our knowledge, levels of STAT3 and STAT5 phosphorylation have not been systematically characterized over the timecourse in which EBV immortalizes primary human cells into LCLs. We therefore infected purified CD19+ peripheral blood B cells with EBV and performed timecourse analysis of EBV latency gene, STAT3 and STAT5 expression, as well as of STAT3 and 5 phosphorylation to indicate their activation status. Interestingly, EBV upregulated STAT3 and STAT5A levels, in particular between days 4 and 21 post-infection, whereas STAT5B levels were relatively constant. Whereas EBV triggered STAT3 phosphorylation, in particular between days 4 and 21 post-infection, a period in which LMP1 levels were markedly elevated and EBNA2 and 3C levels diminished (**Fig. 6A**).

**Figure 6.**
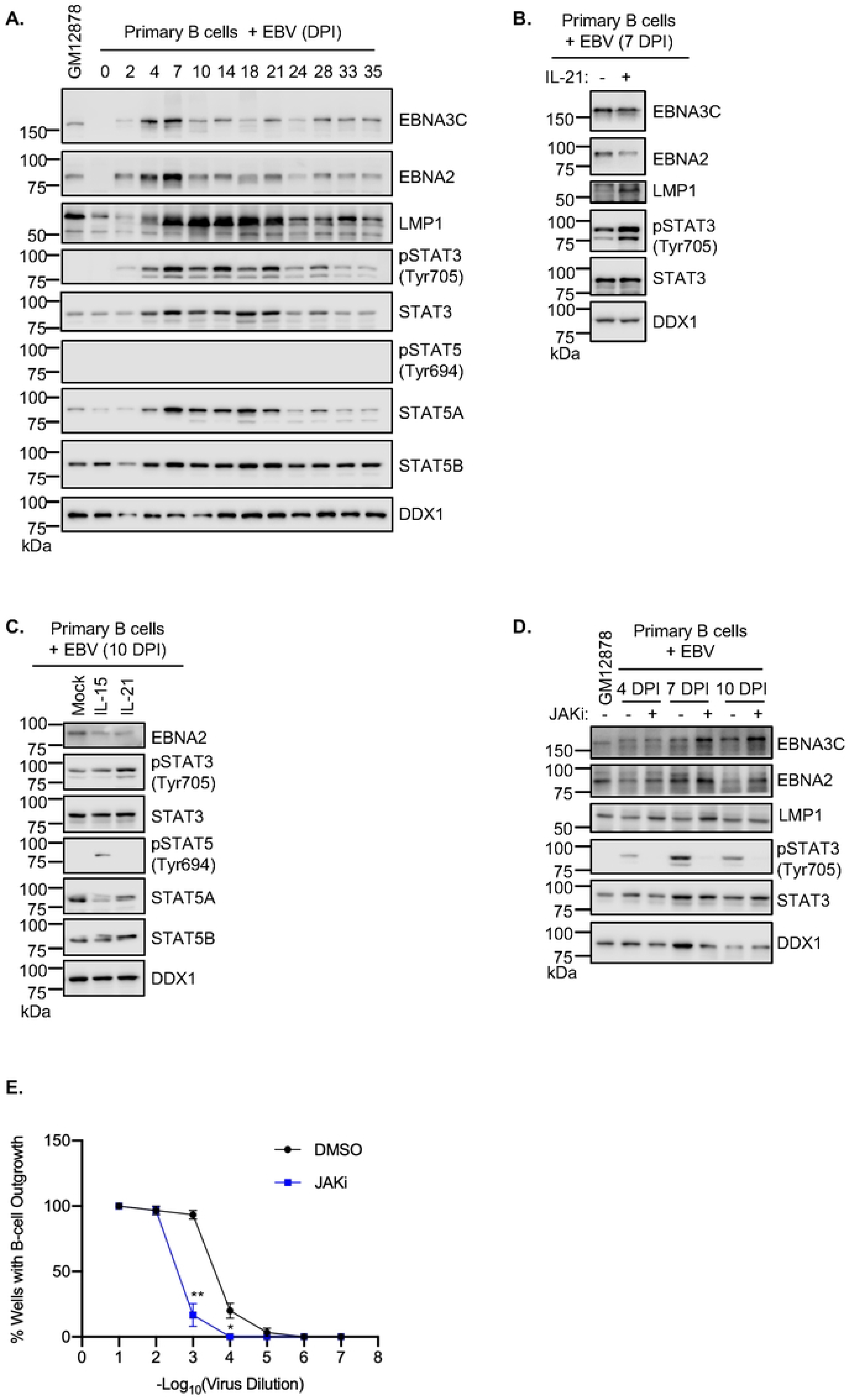
IL-15 and IL-21 remodeling of latency III gene expression in newly infected primary human B cells. (**A**) Immunoblot analysis of WCL from primary human B cells at the indicated days post infection (DPI) by the Akata EBV strain. (**B**) Immunoblot analysis of WCL of primary human B cells at 7 DPI, which were then mock treated or stimulated with IL-21 for six days. (**C**) Immunoblot analysis of WCL from primary B cells at 10 DPI, mock treated or treated with IL-15 or IL-21 for four days. (**D**) Immunoblot analysis of WCL from primary B cells that were treated with DMSO or JAKi (200 ng/ml) for two days at 4, 7 or 10 DPI. GM12878 WCL was included as a control. (**E**) Primary human B-cell transformation assay characterizing effects of DMSO vs JAKi (200 ng/ml) treatment on primary human B-cell outgrowth following infection by Akata EBV. Fitted non-linear regression curves are presented as mean ± SD from n=3 replicates, \**p* < 0.05; \*\**p* < 0.01. Blots are representative of n = 3 replicates. Cytokines were used at 100 ng/ml.

Notably, EBV did not trigger STAT5 phosphorylation, as judged by immunoblot of phosphotyrosine 694 (**Fig. 6A**). We next investigated the effects of IL-21 on EBV latency gene expression when dosed at day 7 post-infection, the earliest timepoint when B-cells begin to convert to lymphoblastoid physiology.^9, 62^ IL-21 reduced EBNA2 and 3C expression and hyper-induced LMP1 (**Fig. 6B**), suggesting conserved STAT roles in EBV latency gene expression in newly infected cells and in LCLs. IL-21 treatment also strongly down-modulated EBNA2 target gene CD23^63, 64^ abundance when applied at multiple timepoints between days 2 and 35 post-infection (**Fig. S9A-B**). We next tested the effects of IL-15 and IL-21 on EBV latency gene expression at day 10 post-infection. Treatment with either cytokine for 4 days reduces EBNA2 expression (**Fig. 6C**).

To characterize the roles of JAK/STAT signaling in EBV-mediated B-cell transformation, we treated newly infected primary human B-cells with JAKi at 4, 7 or 10 DPI. Consistent with JAK/STAT downmodulation of EBNA2 and EBNA3 expression at these early times post-infection, JAKi treatment increased EBNA2 and EBNA3C expression (**Fig. 6D**).

Surprisingly, JAKi treatment also mildly increased LMP1 expression, which likely occurred secondary to increases in EBNA2 levels. JAKi treatment also impaired outgrowth of EBV-infected B-cells in a transformation assay (**Fig. 6E**), suggesting that EBV-driven JAK/STAT signaling supports B-cell immortalization, potentially by titrating the levels of EBV oncoprotein expression. Taken together, our results support a model in which JAK/STAT signaling exerts control over EBV latency gene expression through epigenetic effects on key EBV latency gene promoters (**Fig. 7**).

**Figure 7.**
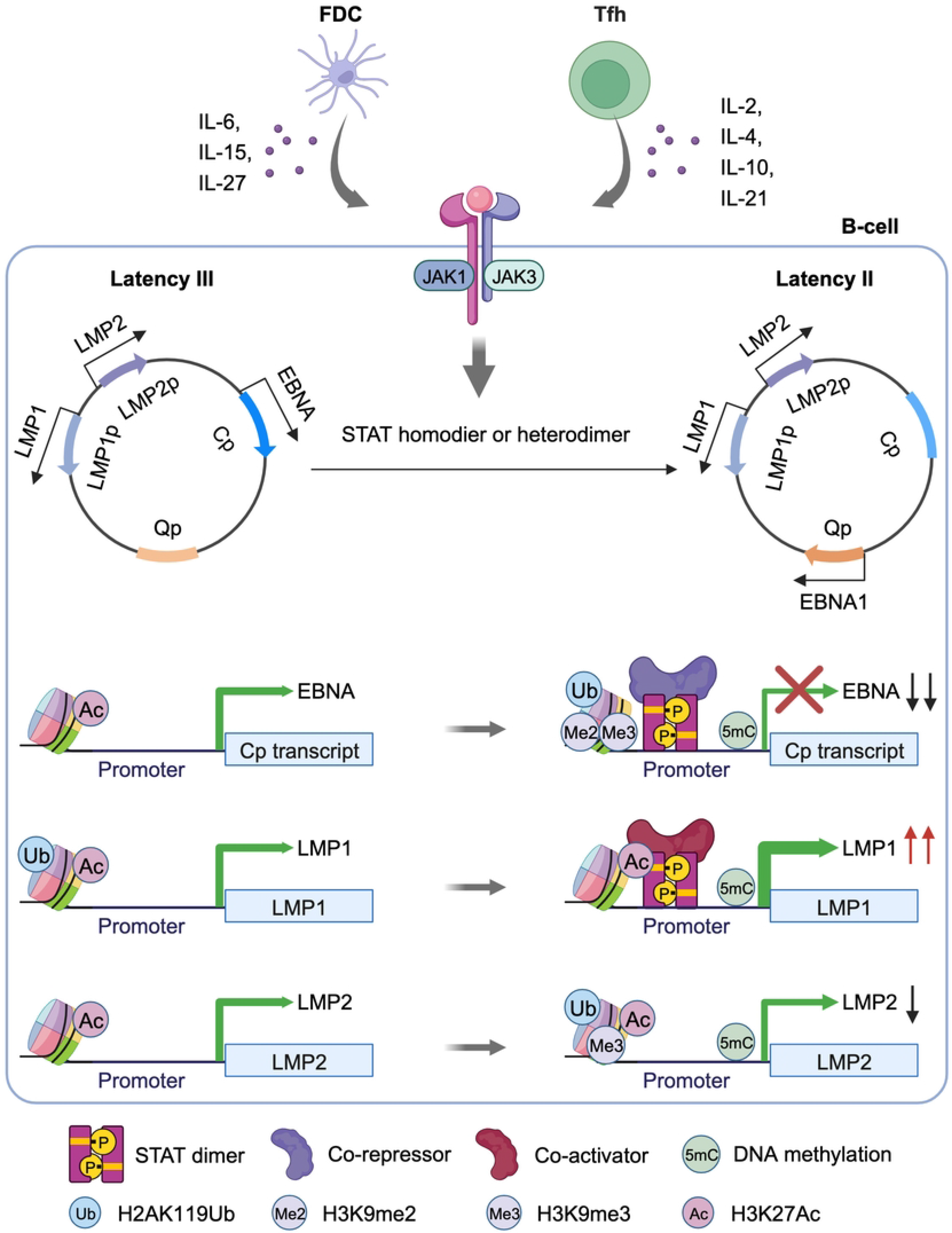
**Model of EBV latency promoter epigenetic remodeling by GC cytokine driven JAK/STAT signaling.**

## Discussion

The EBV germinal center model posits that microenvironmental cues trigger latency program remodeling in order to support infected B-cell survival, immunoevasion and memory B-cell differentiation.^6^ Yet, knowledge has remained incomplete about how specific Tfh and FDC signals alter EBV latency gene promoter epigenomes. Here, we present the first CRISPR analyses to dissect specific STAT roles in latently EBV- infected B-cell responses to GC cytokine cues. We highlight crosstalk between EBV genomic STAT occupancy, histone modification and DNA methylation in GC-cytokine driven reprograming. Our data support a model in which GC cytokines drive a STAT3/5 dependent transcription repressive complex at the EBV genomic C promoter, but instead drive a STAT3 dependent transcription activation complex at the LMP1 promoter (**Fig. 7**). STAT3/5 heterodimers may serve to nucleate a transcription repressive complex at Cp, whereas STAT3 homodimers may instead promote transcription activation at LMP1p. Since STAT3 homodimers support EBNA1 expression in latency II^57, 65^, STAT signaling provides a key means by which EBV- infected cells translate GC microenvironmental cues to the epigenome.

We recently identified that DNA methylation is sufficient for silencing Cp-driven EBNA expression, but that DNA methylation and PRC1 are each important for silencing LMP1 and LMP2A expression in latency I Burkitt cells.^8, 21^ It is therefore noteworthy that IL-21 increased C, LMP1 and LMP2 promoter DNA methylation, but decreased the PRCI H2AK119Ub mark only at the LMP1 promoter in a STAT3/5 dependent manner. IL-21 may alter LMP1 promoter H2AK119Ub abundance by promoting dismissal of PRC1 from LMP1p or instead by recruiting the H2AK119Ub erasers BAP1 or USP16^66^ in a STAT3/5 dependent manner. Importantly, EBNA2 induces and also recruits the TET2 demethylase to the C and LMP promoters.^67, 68^ Therefore, EBNA2 downmodulation by IL-21 likely contributed to the observed increase in EBV genomic methylation.

How then is LMP2A supported in the GC microenvironment upon GC cytokine driven EBNA2 repression? While we studied responses to individual cytokine cues, it is possible that combinatorial signals may be needed to support LMP2A expression.

Alternatively, a distinct GC microenvironmental cue not modelled in our study may be required to support LMP2A expression, such as from dendritic or regulatory T cells.

Thus, a prediction of this model is that on a single cell level, subsets of EBV-infected cells may express LMP1, LMP1 together with LMP2A, or perhaps only LMP2A within distinct GC microenvironmental niches. Such flexibility may support evasion from cytotoxic T-cell responses directed at either LMP1 or LMP2A, may alter the extent of infected cell proliferation or residence time within the GC, and/or may support GC exit upon memory B-cell differentiation into the EBV memory cell reservoir.

Latent EBV infection supports B-cell JAK/STAT signaling, which may provide a basal level to calibrate latency gene expression, even in the absence of Tfh or FDC derived cytokines. The LMP1 C-terminal activation region 3 binds to JAK3^69^, though this region of LMP1 may not by itself be sufficient to activate JAK/STAT signaling.^70, 71^ LMP1 induces IL-10 expression *in vitro*^72^ and together with LMP2A in germinal center B-cells *in vivo*^73^, though levels are likely to be lower than those secreted by Tfh in the GC microenvironment. Furthermore, EBV driven reactive oxygen species accumulation further supports STAT3 activation in the early stages of EBV-driven B-cell outgrowth.^60^ Thus, EBV may have evolved to require a high threshold of JAK/STAT signaling to ensure that latency program selection occurs in the GC microenvironment on the pathway to memory cell differentiation.

While our data suggest that STAT3 is critical for GC cytokine induced remodeling towards latency II, it is noteworthy that IL-21 more strongly induced LMP1 than IL-6, IL- 10 or IL-27, which also strongly activated STAT3. One model to reconcile these observations is that IL-21 signaling may induce a higher abundance of STAT3 homodimers within latency III cells, and these are required for the observed effects on LMP1 expression. Alternatively, IL-21 may more strongly induce co-activators that together with STAT3 upregulate LMP1 expression.

In addition to its roles in EBV latency gene regulation, STAT3 also plays major roles in EBV-driven oncogenic B-cell growth. For instance, B-cells from patients with STAT3 hypomorphic mutation resist EBV-mediated immortalization.^61, 74^ Likewise, transgenic B cell LMP1 expression accelerates lymphomagenesis in a murine model, in which tumors exhibited elevated STAT3 activity.^75^ Elevated STAT3 signaling was also observed in mice with transgenic LMP1 and LMP2A B-cell co-expression.^76^ Relatedly, activated JAK/STAT signaling is observed in EBV+ diffuse large B-cell lymphoma^77^, the Hodgkin lymphoma Reed-Sternberg tumor cell^57^, post-transplant lymphoproliferative disease^78–80^ and plasmablastic lymphoma^81^.

Gamma-herpesviruses may have evolved to subvert STAT3 signaling to support GC- dependent differentiation. EBV, Kaposi’s Sarcoma Associated Herpesvirus and murine gammaherpesvirus 68 (MHV68) have each evolved mechanisms to activate STAT3.^61, 82–87^ STAT3 is important for the establishment of longterm latency by MHV68.^88^ However, in contrast to our findings for EBV, STAT3 does not directly regulate MHV68 viral gene expression, but instead dampens type I IFN responses in newly infected B- cells.^89^ Thus, EBV has evolved specific mechanisms to coopt B-cell STAT signaling to modulate latency gene expression in response to B-cell cues. It is not presently known whether EBV+ B-cells enter GC dark zone structures, in which B-cells undergo multiple rounds of proliferation and somatic hypermutation following stimulation by Tfh and FDC within light zone regions. Since GC cytokine stimulation and STAT3 phosphorylation take place within light zones^90^, it is plausible that EBV+ B-cells may express higher LMP1 levels within light zones, and that they may therefore predominantly reside within GC light zone regions. However, single cell analyses of EBV-infected secondary lymphoid tissue have not yet been performed to address this open area.

In depth understanding of the molecular mechanisms that control EBV latency gene expression may lay the foundation for rational therapeutic approaches. For instance, it may be feasible to target JAK/STAT signaling to downmodulate EBNA expression in tumors that are dependent on the latency III program, such as post-transplant lymphoproliferative disease or central nervous system lymphoma. Conversely, epigenetic approaches to derepress highly immunogenic LMP1 expression may sensitize latency I tumors such as Burkitt lymphoma to antiviral T-cell surveillance, including adoptive transfer of T-cells reactive with LMP1 derived epitopes.^91, 92^ Furthermore, LMP1 de-repression promises to re-sensitize EBV-infected latency I tumors to T-cell responses to tumor associated antigens.^93^

In summary, multiple FDC and Tfh derived cytokines repress Cp driven EBNA expression, whereas IL-21 and to a lesser extent IL-4 and IL-10 support LMP1 expression through STAT dependent EBV epigenomic remodeling. STAT3 and 5 were critical for cytokine mediated Cp silencing, whereas STAT3 was critical for LMP1 hyper-induction. GC cytokine signaling increased repressive epigenetic marks, including DNA methylation and H2AK119Ub, while decreased active chromatin mark H3K27Ac at Cp in EBV latency III cells. However, IL-21 increased H3K27Ac at the LMP promoters, but decreased H2AK119Ub only at the LMP1 promoter in both EBV latency III and I cells.

IL-21 also decreased DNA methylation at LMP1 promoter in EBV latency I Burkitt cells. Therefore, STAT3 and 5 serve as major hubs of EBV epigenomic remodeling in response to GC cytokine signaling to support latency program remodeling.

## Acknowledgements

This work was supported by NIH U01CA275301, R01 AI164709, R21 AI170751, by a Lymphoma Research Foundation Postdoctoral Fellowship to Y.L. and by T32 T32AI007245 to N.B.

## Data Availability Statement

All RNA-seq datasets will be released in the NIH GEO omnibus upon publication.

## Author contributions

Y.L. and J.Y. performed the experiments and data analysis; RNA-seq bioinformatics analysis was performed by N.B.; Y.L., L.G.R., E.C. and B.E.G. supervised the study. Y.L., J.Y., N.B., and B.E.G. wrote the manuscript. Y.L. and J.Y. contributed equally to this work.

## Materials and Methods

### Cell culture

EBV+ latency I cells, P3HR1 (A gift from Dr. Elliott Kieff), Akata (A gift from Dr. Elliott Kieff), Mutu I (A gift from Dr. Jeffrey Sample) and Kem I (A gift from Dr. Jeffrey Sample), and latency III cells, GM12878 (purchased from Coriell Institute), GM12881 (purchased from Coriell Institute), Kem III (A gift from Dr. Jeffrey Sample) and Jijoye (purchased from American Type Culture Collection, ATCC), were all grown in Roswell Park Memorial Institute (RPMI) 1640 medium with 10% fetal bovine serum (FBS). 293T cells (purchased from ATCC) were grown in Dulbecco’s Modified Eagle’s Medium (DMEM) with 10% FBS. All B cell lines used in this study stably express *Streptococcus pyogenes* Cas9, which were generated by lentiviral transduction followed by blasticidin selection.^94^ All cells were grown in a humidified chamber with 5% carbon dioxide at 37°C.

### Cytokines and JAKi treatment

Latency I and III B cells were seeded at 500,000 cells/ml in 12-well plates and mock treated with PBS or cytokines (**Table S3**) at 50 ng/ml and 100 ng/ml, respectively. EBV infected human primary B cells were treated with IL-15 or IL-21 at 100 ng/ml. For long-term treatment, cells were re-seeded with fresh culture medium supplemented with indicated cytokines, which were refreshed every 48 hours. For JAK inhibitor treatment, cells were pre-treated with JAK inhibitor I (JAKi) at indicated doses for one hour at 37 °C, followed by cytokine treatment, refreshed every 48 hours.

### CRISPR/Cas9 editing

CRISPR/Cas9 editing was performed as previously described.^95, 96^ In brief, Brunello library^97^ single guide RNA (sgRNA) were cloned into pLentiGuide-puro (a gift from Feng Zhang, Addgene plasmid #52963), pLenti-spBsmBI-sgRNA-Hygro (a gift from Rene Maehr, Addgene plasmid #62205), or pLentiGuide-zeo (a gift from Rizwan Haq, Addgene plasmid #160091). sgRNA sequences were verified by Sanger sequencing. All sgRNAs used in this study are listed in **Table S4**. Target cells were transduced with lentivirus expressing sgRNAs against the target gene, or as a control, against GFP (pXPR-011, a gift from John Doench). Lentivirus were produced by transfection of 293T cells with pCMV-VSV-G (a gift from Bob Weinberg, Addgene plasmid #8454), psPAX2 (a gift from Didier Trono, Addgene plasmid #12260), and the sgRNA expression vector using the TransIT-LT1 transfection reagent. 293 supernatants were added to target B-cells at 48 and 72 hours post-293 transfection. Transduced cells were then selected with puromycin (3 μg/ml) for 3 days.

For STAT5A and STAT5B combinatorial editing, GM12878 or Jijoye Cas9+ cells were initially transduced with lentivirus expressing STAT5A sgRNA and selected with hygromycin (200 μg/ml) for 7 days, followed by transduction with lentiviruses expressing STAT5B sgRNA and selected with puromycin (3 μg/ml) for 3 days. For experiments with STAT5A/5B/3 editing, STAT3 was depleted in STAT5A/5B edited GM12878 cells by transduction with lentivirus expressing STAT3 sgRNA, and transduced cells were selected by zeomycin (200 μg/ml) for 7 days. On-target CRISPR effects were validated by immunoblotting.

### cDNA cloning and transduction

cDNA entry vectors used in this study are listed in **Table S4**, which were purchased from DNASU and Addgene. STAT1, 3, and 6 cDNA were sub-cloned into the destination vector pLX-TRC313 (a gift from John Doench) and STAT3_p.A662C_N664C (constitutively active STAT3 with A662C and N664C mutations)^44^ was cloned into pLIX- 402 (a gift from John Doench) by Gateway LR recombination. As described previously^98^, the destination vector and donor vector containing the gene of interest were co-incubated with 1x LR Clonase Enzyme Mix overnight at room temperature. The reaction mixture was then transformed into Stbl3 competent cells and plated on LB agar plate containing ampicillin. Destination vectors were used to make lentiviruses, which were used to transduce target B-cells. Transduced cells were selected by puromycin or hygromycin for pLIX-402 or pLX-TRC313 vectors, respectively.

### Immunoblotting

Immunoblotting analysis was performed as previously described^21^, Cells were lysed in 1x Laemmli Sample Buffer and sonicated briefly. For detection of LMP2A, cells were lysed with M-PER™ Mammalian Protein Extraction Reagent and incubated on ice for 30 minutes. Lysates were centrifuged at 15,000 x g for 15 minutes. 2x Laemmli Sample Buffer was added into supernatant and boiled at 70 °C for 10 minutes. Lysates were resolved by SDS-PAGE and transferred onto nitrocellulose membranes, which were blocked with 5% nonfat milk in TBST buffer for 1 hour and then incubated with primary antibodies at 4 °C overnight. Blots were then washed 3 times with TBST, followed by secondary antibody incubation for 1 hour at room temperature. Blots were washed 3 times in TBST buffer and were developed with the ECL chemiluminescence substrate. Images were captured by a LI-COR Fc platform. All antibodies used in this study are listed in **Table S3**.

### Flow cytometry assay

Cells were washed once with FACS buffer (2% FBS v/v, PBS), followed by incubation with primary antibodies in FACS buffer for 30 minutes at room temperature in the dark. Labeled cells were palleted, washed twice and resuspended in FACS buffer into flow cytometry-compatible tubes and processed immediately. Flow cytometry data was recorded with a BD FACSCalibur instrument and analyzed with FlowJo X software.

### Akata virus production, primary B cells isolation and infection

EBV was produced from EBV+ Akata cells. In brief, EBV+ Akata cells were resuspended in FBS-free RPMI media at 2-3 million cells/mL and induced with 0.25% (v/v) goat anti-human immunoglobulin G serum for 6 hours at 37 °C. Cells were then pelleted and resuspended in 4% FBS RPMI media and cultured in 37 °C for 3 days. Supernatant were then collected and filtered through 0.45 µM filter. Viruses were 50-fold concentrated by ultracentrifugation and stored at -80 °C until use.

Primary B-cells were isolated by negative selection from discarded, de-identified peripheral blood mononuclear cells from the Brigham and Women’s Hospital Blood Bank, obtained following platelet donation, using an Institutional Review Board approved protocol and donor informed consent. RosetteSep and EasySep negative isolation kits were used according to the manufacturer’s instructions to isolate CD19+ B-cells. B cells were then cultured with RPMI containing 10% FBS. Primary B cells were seeded at 500,000 cells/ml and infected by the Akata EBV strain at multiplicity of infection (MOI) of 0.1, as determined by the Green Daudi assay.

### Primary human B cell EBV transformation assay

EBV transformation assays were performed as described previously.^99^ Briefly, purified human primary B cells were infected with Akata EBV using serial 10-fold dilutions. Cells were cultured with media containing DMSO or JAKi (200 ng/ml) and were seeded in 96- wells plates at 500,000 cells/ml (30 wells per condition). Media containing DMSO or JAKi was refreshed every three to four days. The percentage of wells positive for B-cell outgrowth at four weeks post infection was calculated and plotted relative to the dilution of virus.

### Chromatin immunoprecipitation (ChIP) assay

After cytokine treatment, 10 million cells were cross-linked with 1% formaldehyde in 10 ml growth medium for 10 minutes, followed by quenching with 2.5M glycine in distilled water for 5 minutes. Cells were washed with ice-cold PBS three times and then lysed in 0.5 ml 1% SDS lysis buffer (50 mM Tris, 10 mM EDTA, 1% SDS), supplemented with 1x cOmplete™, EDTA-free Protease Inhibitor Cocktail. Chromatin was fragmented using a Bioruptor Pico sonication device with 30s on/ 30s off (20 cycles for GM12878 cells, 12 cycles for Mutu I cells), and centrifuged at 13,200 rpm for 10 mins at 4 °C. This protocol resulted in fragments of average length 100-200 bp, to enable differentiation of STAT occupancy at closely spaced EBV genomic STAT binding sites. Supernatants were removed and then diluted 1:10 in ChIP dilution buffer (1.2 mM EDTA, 16.7 mM Tris, 167 mM NaCl, 0.01% SDS, 1.1% Triton X-100) supplemented with protease inhibitor cocktail. Chromatin from one million cells was used for each ChIP reaction. 1% of sonicated chromatin was saved as input and stored at -80 °C until use. Diluted chromatin was rotated overnight at 4 °C with the indicated antibody and 20 µl protein A+G magnetic beads. Next day, beads were pelleted, washed twice with a lower salt buffer (150 mM NaCl, 2 mM EDTA, 20 mM Tris, 0.1% SDS, 1% Triton X-100) and then a high-salt buffer (500 mM NaCl, 2 mM EDTA, 20 mM Tris, 0.1% SDS, 1% Triton X-100), and once with LiCl buffer (0.25 M LiCl, 1% NP-40, 1% sodium deoxycholate, 1 mM EDTA, 10 mM Tris) and finally TE buffer (10 mM Tris, 1 mM EDTA). Chromatin was eluted in Elution buffer (100 mM NaHCO3, 1% SDS) and reverse cross-linked at 65 °C for 2 hours. QIAquick PCR purification kits were used to purify the immunoprecipitated DNA, followed by qPCR with PowerUp SYBR green PCR master mix on a CFX Connect Real-Time PCR Detection System (Bio-Rad). All reagents, antibodies and primers used for ChIP are listed in **Tables S3 and S4**.

### Methylated DNA immunoprecipitation (MeDIP) assay

Genomic DNA was extracted using DNeasy Blood& Tissue Kit, followed with MeDIP assay with MagMeDIP kit, following the manufacturer’s protocol. qPCR was then performed with primers specifically target EBV promoters. All reagents and primers used for MeDIP are listed in **Tables S3 and S4**.

### RNA-seq and data analysis

mRNA was isolated via the RNeasy Mini kit with in-column genomic DNA digestion protocol was followed, according to the manufacturer’s instructions. To construct indexed libraries, 1 μg of total RNA was used for polyA mRNA purification, using the NEBNext Poly(A) mRNA Magnetic Isolation Module, followed by library preparation using the NEBNext Ultra RNA Library Prep with Sample Purification Beads. Each experimental treatment was performed in biological triplicate. Libraries were multi-indexed, pooled and sequenced on an Illumina NovaSeq 6000 using PE150 Sequencing Strategy by Novogene Corporation. Adaptor-trimmed reads were mapped to Akata EBV genome (Accession#: KC207813.1) or human GRCh37.83 transcriptome assembly using salmon (v1.10.0). Quality control was performed using fastqc.

Differentially expressed genes were identified in R (v4.0.3) using DESeq2^100^ under default settings with the apeglm shrinkage estimator (https://doi.org/10.1093/bioinformatics/bty895) and annotations derived from the hg19 build from Ensembl release 75^101^ and accessed via biomaRt. Volcano plots were generated in GraphPad Prism 8, using Log2 (Fold Change) and −Log10 (p value) data. Differentially expressed genes from each condition were subjected to Enrichr analysis and top 10 KEGG pathways with adjusted *p* value < 0.05 cutoff were visualized. All reagents and kits used for RNA-seq are listed in **Table S3**.

### Quantification and statistical analysis

All immunoblots were performed with three independent experiments and qPCR was performed in three independent experiments. Statistical significance was assessed with Student’s t test using GraphPad Prism 8 software, where NS = not significant, *p* > 0.05; * *p* < 0.05; ** *p* < 0.01; *** *p* < 0.001. Biorender was used to create the schematic models.

## Supplementary Figure Legends

Figure S1. **GC cytokine effects on latency III B-cell EBV and host gene expression.** (**A**) Schematic of Tfh and FDC cytokine driven JAK/STAT signaling. (**B-C**) Immunoblot analysis of WCL from GM12881 (B) and latency III Jijoye (C) cells treated with the indicated cytokines for six days. (**D**) Immunoblot analysis of WCL from GM12878 and Kem III cells six days post mock, IL-15 or IL-21 treatment. (**E**) Volcano plot (left) and KEGG pathway analysis (right) of host genes expression in GM12878 stimulated by IL-15 versus mock-simulated for six days from n = 3 independent replicates. The top 10 most differentially expressed KEGG pathways are shown. (**F**) Volcano plot (left) and KEGG pathway analysis (right) of host genes expression in GM12878 stimulated by IL-21 versus mock-simulated for six days from n = 3 independent replicates. Cytokines were used at 100 ng/ml and were refreshed every two days. Immunoblots are representative of n = 3 replicates.

Figure S2. GC cytokine effects on latency I B cell EBV and host gene expression.

(**A**) Immunoblot analysis of WCL from latency I Kem I Burkitt B cells treated with the indicated cytokines for 24 hours. (**B**) Immunoblot analysis of WCL from Mutu I and Kem I treated with IL-21 for one or two days, as indicated. (**C**) Immunoblot analysis of WCL from Mutu I and Kem I one day post mock, IL-10 or IL-21 treatment. GM12878 WCL was included as a positive control. (**D**) Volcano plot (left) and KEGG pathway analysis (right) of differentially expressed Mutu I host genes one day after IL-4+CD40L vs mock stimulation from n=3 independent replicates. (D) Volcano plot (left) and KEGG pathway analysis (right) of differentially expressed Mutu I host genes one day after IL-4+CD40L vs mock stimulation from n=3 independent replicates. The top 10 KEGG pathways amongst differentially regulated genes are shown. Cytokines and CD40L were used at 50 ng/ml for EBV latency I cells. Immunoblots are representative of n = 3 replicates.

Figure S3. **STAT3 and 5 roles in IL-15 and IL-21 driven EBV latency III gene regulation.** (**A**) Immunoblot analysis of WCL from latency III Jijoye B cells expressing control sgRNA or sgRNA targeting STAT5A and STAT5B, mock treated or treated with IL-15 or IL-21 for six days. (**B-C**) Immunoblot analysis of WCL from Jijoye (B) or GM12878 (C) expressing control sgRNA or sgRNA targeting the indicated STAT transcription factor gene, mock treated or treated with IL-10 or IL-21 for six days. Blots are representative of n = 3 replicates. Cytokines were used at 100 ng/ml and refreshed every 2 days.

Figure S4. IL-21 effects on LCL EBNA2 and LMP1 expression are not dependent on BCL6 but correlate with STAT-dependent LMP promoter methylation. (A) Immunoblot analysis of WCL from GM12878 cells expressing control sgRNA or independent *BCL6* targeting sgRNA that were mock treated or treated with IL-21 (100 ng/ml) for two or four days. Blot is representative of n = 3 replicates. (B) Flow cytometry analysis of LMP1 target ICAM-1 and EBNA2 target CD300A plasma membrane expression in GM12878 expressing control sgRNA or BCL6 sgRNA and mock treated or IL-21 treated for 2 or 4 days, as indicated. (C) MeDIP-qPCR analysis of GM12878 expressing control or sgRNA targeting STAT3/5A/5B, mock treated or treated with IL-15 or IL-21 for six days, followed by qPCR with primers targeting the LMP1 promoter (LMP1p, left) or LMP2 promoter (LMP2p, right). Mean ± SD ChIP-qPCR % input values from n = 3 replicates are shown. \**p* < 0.05; \*\**p* < 0.01; \*\*\**p* < 0.001.

Figure S5. **STAT roles in LMP1 de-repression in GC cytokine treated latency I B cells.** (**A**) Immunoblot analysis of WCL from Mutu I cells treated with JAKi (0-1,000 ng/ml) for one hour, followed by IL-21 treatment for one or two days. GM12878 cell lysate was included as a positive control. (**B**) Immunoblot analysis of WCL from Kem I expressing control sgRNA or sgRNA targeting STAT1 or STAT3, mock treated or treated with the indicated cytokine for one day. (**C**) Immunoblot analysis of WCL from Mutu I conditionally induced for control GFP or constitutively active STAT3 for one day by 0.5 or 1 µg/ml doxycycline. (**D**) Immunoblot analysis of Mutu I expressing the indicated control GFP or STAT cDNA and stimulated as indicated for 1 day. (**E**) MeDIP-qPCR of the LMP2 promoter (left) and C promoter (right) in Mutu I and Kem I, mock treated or IL- 21 treated for one day. Mean ± SD input % of n = 3 replicates are shown, \**p* < 0.05; \*\**p* < 0.01. All cytokines were used at 50 ng/ml. Blots are representative of n = 3 replicates.

Figure S6. **STAT roles in IL-15 and IL-21 driven LMP1 and LMP2 promoter epigenetic remodeling.** (**A-C**) ChIP-qPCR analysis of LMP1 promoter H3K9me2 (A), H3K9me3 (B) or H3K27me3 (C) abundances from GM12878 expressing control or STAT3/5A/5B targeting sgRNAs, mock treated or treated with 100ng/ml IL-15 or IL-21 for six days. (**D-F**) ChIP-qPCR analysis of LMP2 promoter H3K9me2 (D), H3K9me3 (E) or H3K27me3 (F) abundances in GM12878 expressing control or STAT3/5A/5B targeting sgRNAs, mock treated or treated with IL-15 or IL-21 for six days. Mean ± SD input % of n = 3 replicates are shown, \**p* < 0.05; \*\**p* < 0.01.

Figure S7. **STAT3 roles in LMP1 and LMP2 promoter IL-21 driven epigenetic remodeling in latency I B-cells.** (**A-B**) ChIP-qPCR analysis of LMP1 promoter H3K9me2 (A) or H3K9me3 and H3K27me3 (B) abundances from Mutu I expressing control or STAT3 targeting sgRNA, mock treated or treated with IL-21. (**C-F**) ChIP-qPCR analysis of LMP2 promoter H3K27Ac (C) or H2AK119Ub (D), H3K9me2 (E) or H3K9me3 and H3K27me3 (F) abundances in Mutu I expressing control or STAT3 targeting sgRNAs, mock treated or treated with IL-21. Cells were treated with 50 ng/ml IL-21 for one day. Mean ± SD input % of n = 3 replicates are shown, \**p* < 0.05; \*\**p* < 0.01.

Figure S8. **STAT3 roles in IL-15 and IL-21 driven Cp epigenetic remodeling.** (**A**) ChIP-qPCR analysis of Cp H3K27me3 abundances in Mutu I expressing control or STAT3/5A/5B targeting sgRNAs, mock treated or treated with IL-15 or IL-21 (100ng/ml) for six days. (**B-F**) ChIP-qPCR analysis of Cp H3K27Ac (B), H2AK119Ub (C), H3K9me2 (D), H3K9me3 (E) or H3K27me3 (F) abundances in Mutu I expressing control or STAT3 targeting sgRNAs, mock treated or treated with IL-21 50 ng/ml for 1 day. Mean ± SD input % of n = 3 replicates are shown, \**p* < 0.05; \*\**p* < 0.01.

Figure S9. **IL-21 effects on newly EBV infected primary B-cell EBNA2 target gene CD23 expression.** (**A**) Plasma membrane CD23 abundances in primary human B-cell mock treated or treated with IL-21 (100 ng/ml) at Day 7 vs 18 post-infection by Akata EBV. IL-21 was refreshed every 2 days. (**B**) Mean ± SD CD23 abundances from n = 3 replicates of primary B-cells infected by Akata EBV in the absence or presence of IL-21, as in (A), \*\*\**p* < 0.001.

## Supplementary Tables

Table S1. RNA-seq of EBV Gene Expression

Table S2.RNA-seq of Host Gene Expression

Table S3. Reagents, Antibodies and Kits

Table S4. sgRNAs, plasmids and primers

